# Myocardial IGF2R is a critical mediator of inflammation and fibrosis after ischemia-reperfusion injury

**DOI:** 10.1101/2023.04.21.537835

**Authors:** Zhichao Wu, Wei Huang, Xingyu He, Suchandrima Dutta, Christian Paul, Guo-Chang Fan, Onur Kanisicak, Meifeng Xu, Jialiang Liang, Yigang Wang

**Affiliations:** Department of Pathology and Laboratory Medicine, College of Medicine, University of Cincinnati, Cincinnati, OH, USA, 45267; Department of Internal Medicine, College of Medicine, University of Cincinnati, Cincinnati, OH 45267; Department of Pharmacology and Systems Physiology, College of Medicine, University of Cincinnati, Cincinnati, OH, USA, 45267

## Abstract

Ischemia-reperfusion (I/R) injury is a common occurrence in various surgical procedures used to treat heart diseases. However, the role of insulin-like growth factor 2 receptor (IGF2R) during the process of myocardial I/R remains unclear. Therefore, this study aims to investigate the expression, distribution, and functionality of IGF2R in various I/R-associated models (such as reoxygenation, revascularization, and heart transplant). Loss-of-function studies (including myocardial conditional knockout and CRISPR interference) were performed to clarify the role of IGF2R in I/R injuries. Following hypoxia, IGF2R expression increased, but this effect was reversed upon restoration of oxygen levels. Loss of myocardial IGF2R was found to enhance the cardiac contractile functions, and reduced cell infiltration or cardiac fibrosis of I/R mouse models compared to the genotype control. CRISPR-inhibition of IGF2R decreased cell apoptotic death under hypoxia. RNA sequencing analysis indicated that myocardial IGF2R played a critical role in regulating the inflammatory response, innate immune response, and apoptotic process following I/R. Integrated analysis of the mRNA profiling, pulldown assays, and mass spectrometry identified granulocyte-specific factors as potential targets of myocardial IGF2R in the injured heart. In conclusion, myocardial IGF2R emerges as a promising therapeutic target to ameliorate inflammation or fibrosis following I/R injuries.

## INTRODUCTION

Ischemia-reperfusion (I/R) injury is a well-known phenomenon that occurs in various surgical procedures to treat heart diseases, such as percutaneous coronary intervention for acute myocardial infarction, heart transplantation, and coronary artery bypass grafting.^1^ While several pharmacological interventions have been investigated to improve I/R injury, no single therapy can effectively target the multiple pathways involved in this complex process.^2^ Eventually, the activation of irreversible cell death pathways after I/R can contribute to the development of heart failure. Understanding the molecular and cellular mechanisms underlying I/R injury is critical for refining the existing surgical approaches to prevent or mitigate their adverse effects on the heart.

Currently, most of the drugs being evaluated for their cardioprotective effects are derived from the cardioprotective signal transduction pathways.^3, 4^ The activation of prosurvival pathways (such as PI3K/AKT, JAK/STAT, and ERK) modulated by the insulin-like growth factor (IGF) system is essential for cardiac homeostasis including cell growth or death and regulation of energy metabolism.^5-7^ Previous studies have established the homology between insulin, IGF1, and IGF2, as well as elucidated the interactions among the insulin receptor (INSR), IGF1 receptor (IGF1R), and IGF2 receptor (IGF2R).^8-10^ As the administration of insulin, IGF1, IGF2, and other growth factors gains increasing interest as a potential treatment for myocardial I/R,^11-13^ investigating the receptors of these factors could provide valuable insights into the role of the IGF system in cardioprotection.

It is well-known that canonical downstream of IGF1R such as PI3K/AKT can maintain outer mitochondrial membrane potential and inhibit glycogen synthase kinase-3β, thereby protecting cells against necrosis triggered by I/R.^14-16^ Recent research has shown that inhibiting IGF2R signaling promotes cardiomyocyte proliferation and reduces hypoxia- or TNFα-induced cell death,^17-19^ while the role of IGF2R signaling in I/R remains largely unknown. With broad expression across many cell types, IGF2R is a versatile receptor that can bind multiple ligands (including IGF2 and M6P-modified proteins) and plays a critical role in intracellular transport, shuttling between the trans-Golgi network, endosomes, and plasma membrane.^20, 21^ Therefore, disruption of the Golgi-to-lysosome transport by loss of IGF2R can lead to reduced lysosomal activity, resulting in the accumulation of misfolded proteins and increased production of reactive oxygen species due to dysfunction in both autophagy and mitophagy pathways.^22, 23^ Given its multifunctional nature, the effects of IGF2R on cell viability appear to be cell type-specific and context-dependent under stress conditions. In this study, the loss-of-function models were used to investigate the role of IGF2R in myocardial I/R injuries.

## RESULTS

### Dynamic expression of IGF2R during myocardial I/R

The expression of IGF2R in mouse hearts exhibited an initial increase following myocardial ischemia (MI) and reperfusion as compared to the sham operation but subsequently decreased to baseline levels after a 6-hour reperfusion period (**Fig.1A**). In addition to cardiomyocytes, IGF2R was expressed in non-CM cells located in the interstitial space of reperfused heart (**Fig.1B**). The single-cell RNA-seq dataset (Tabula Muris)^24^ also provides evidence that IGF2R is expressed constitutively across multiple cell types within the heart (**Fig.1C**). To investigate the precise role of myocardial IGF2R in I/R, we generated conditional knockout (cKO) models by crossing Cre-inducible *Igf2r*^*fl/fl*^ mice with *Myh6*^*MerCreMer*^ mice and treating the resultant offspring with tamoxifen (**Fig.1D**), while *Igf2r*^*fl/fl*^ litters were used as the control group (Ctrl). IGF2R gene expression was significantly decreased in cKO as confirmed by PCR and western blotting (**Fig.1E-F**).

**Fig.1.**
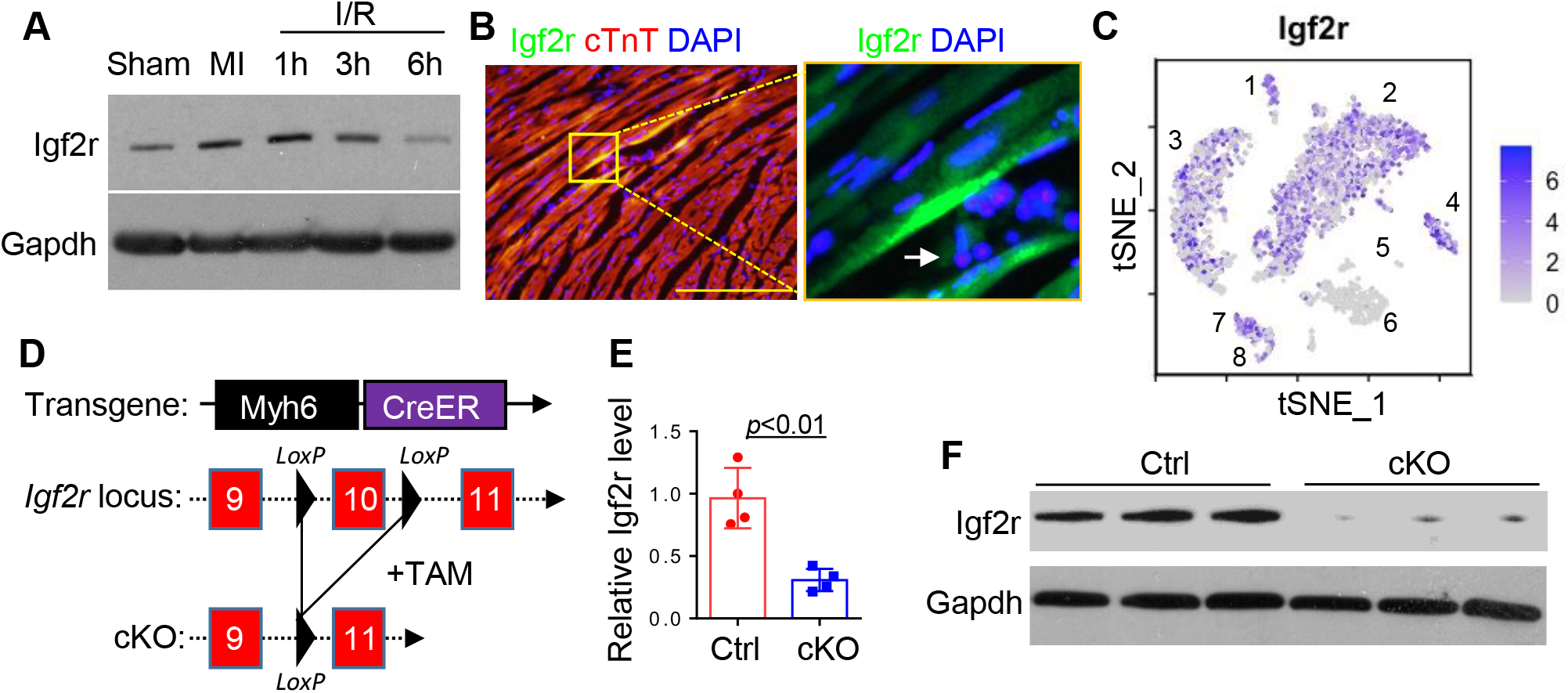
IGF2R expression in ischemic or reperfused hearts. (**A**) Immunoblotting of IGF2R and GAPDH (loading control) in mice after 45min myocardial ischemia (MI) induced by left anterior descending artery ligation or releasing ligature (I/R, ischemia-reperfusion) for 1 to 6 hours. (**B**) Immunostaining of IGF2R and cTnT in the ischemic heart reperfused for 1 hour. Scale bar, 50μm. Arrow indicates noncardiac cells. (**C**) Single-cell profiling of IGF2R expression in normal mouse heart (reanalysis of GSM2967050). Cell phenotype annotation: 1-cardiac muscle cell; 2-fibroblast; 3-endothelial cell; 4-Endocardial cell; 5-cardiac neuron; 6-leukocyte; 7-myofibroblast; 8-smooth muscle cell. (**D**) Schematic of transgenic mouse models for conditional knockout (cKO) of myocardial IGF2R. (**E**) Expression of IGF2R mRNA in the Ctrl (Igf2r^fl/fl^) or cKO (*Myh6*^*MerCreMer*^*;Igf2r*^*fl/fl*^) hearts after the tamoxifen (TAM) diet for 4 weeks. (**F**) Expression of IGF2R protein in the Ctrl or cKO hearts after the TAM diet for 4 weeks.

### Loss of myocardial IGF2R improves heart function of I/R models

Subsequently, two different types of IR injury were employed to assess the effect of cKO on heart function. In a form of warm I/R (**Fig.2A**), the heart function of Ctrl or cKO mice was analyzed by echocardiography after LAD ligation surgery. At 1 week after I/R, there was no difference in the ejection fraction (EF) between Ctrl and cKO (**Fig.2B**). The heart function of cKO mice demonstrated an improvement at 4 weeks post-I/R when compared to Ctrl mice, with increases in EF, fractional shortening, and interventricular septal (**Fig.2C-D**). In addition, heterotopic cardiac transplant models were established to mimic cold I/R injury (**Fig.2E**). Following transplantation within 12-hour preservation in cold HTK solutions, there was no difference in re-beating time between the control group and the cKO donor hearts (**Fig. 2F**). Although the re-beating time of cKO hearts was shorter than that of the control group after 24-hour preservation in cold HTK solutions, both grafts had a dramatically decreased median survival time (**Fig. 2G**).

**Fig.2.**
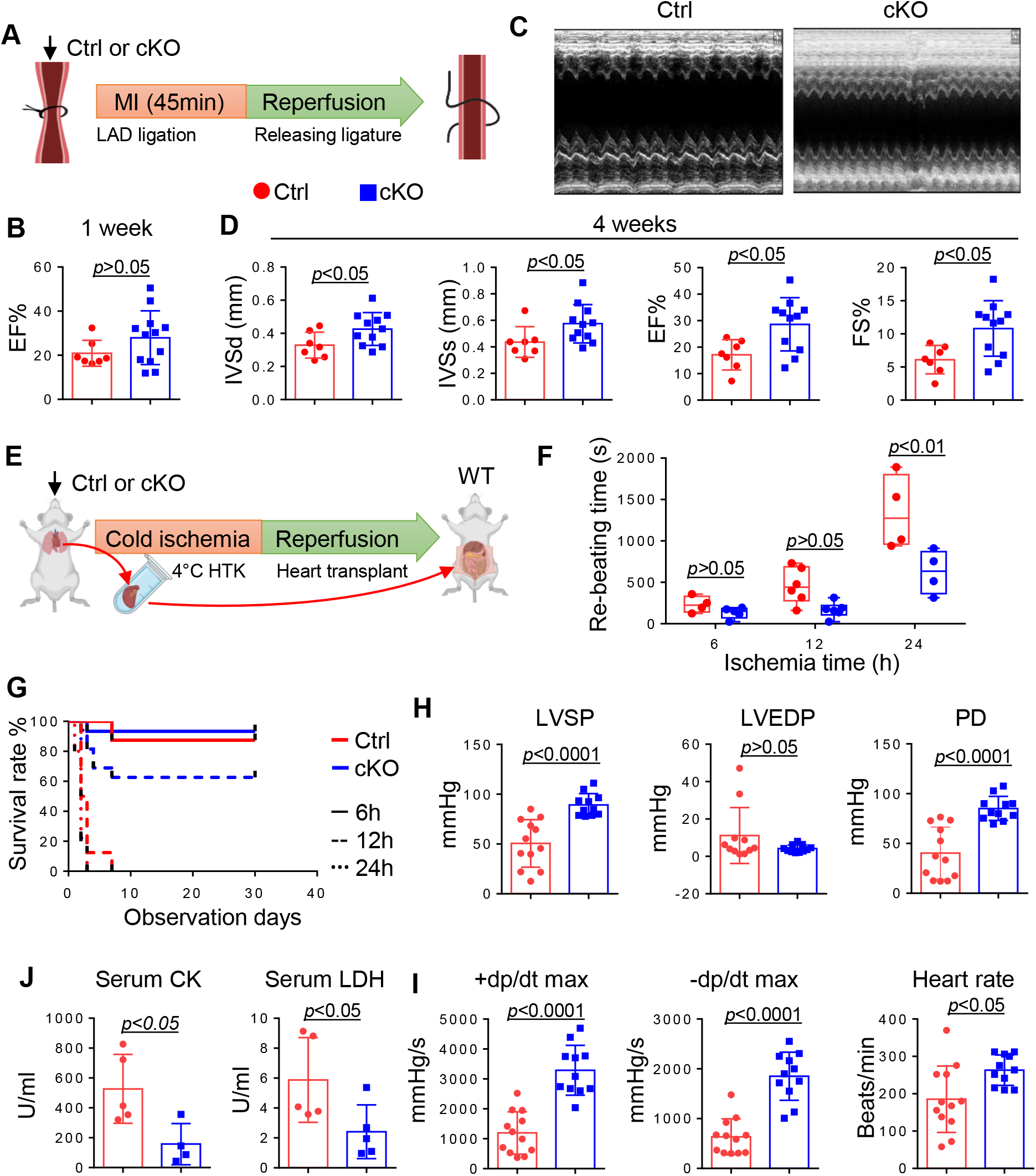
IGF2R-cKO enhances cardiac contractile functions. (**A**) Schematic of the warm I/R model. Myocardial ischemia (MI) induced by left anterior descending artery (LAD) ligation for 45min and reperfusion by releasing the ligature. (**B**) Heart function of control and cKO mice assessed by echocardiography in 1 week after the warm I/R. (**C-D**) Heart function of control and cKO mice assessed by echocardiography in 4 weeks after the warm I/R. EF-ejection fraction; FS-fractional shortening; IVSd-interventricular septal end-diastole; IVSs-interventricular septal end-systole. (**E**) Schematic of the cold I/R model. Collecting and perfusing donor hearts with 4°C HTK solution and keeping on ice until transplant into the abdomen of syngeneic recipients. (**F**) The rebeating time (after anastomosis) for ventricular contraction restoration of control or cKO heart grafts (preserved for 6, 12, or 24 hours in cold HTK). (**G**) Survival curves of control or cKO heart grafts after transplant. The complete cessation of a heartbeat is the endpoint. (**H-I**) Heart rate, pressure difference (PD), left ventricle systolic pressure (LVSP), left ventricle end-diastolic pressure (LVEDP), maximum dp/dt in the systolic period, and maximum dp/dt in the diastolic period of control or cKO heart grafts (preserved for 12 hours in HTK) on day 1 post-transplant. (**J**) The serum creatine kinase (CK) or lactate dehydrogenase (LDH) levels of control or cKO heart grafts (preserved for 12 hours in HTK) on day 1 post-transplant.

All Ctrl and cKO donor hearts survived after 6-hour preservation without a significant difference, but ∼60% of cKO hearts exhibited a significantly longer median survival time than Ctrl hearts after 12-hour preservation (**Fig.2G**), suggesting a potential protective effect of cKO against ischemic injury. Therefore, on day one post-surgery, a catheter-based approach was used to evaluate the left ventricular (LV) pressure and contractility of Ctrl or cKO hearts that underwent 12-hour preservation (**Fig.2H-I**). The LV systolic pressure and heart rate were increased in cKO donor hearts as compared with the control, while there was no significant difference in the LV end-diastolic pressures. The +dp/dt max and the -dp/dt max were rapidly raised in cKO donor hearts, reflecting an improved contractile function. Furthermore, recipients with cKO hearts demonstrated a significant decrease in serum levels of intracellular enzymes, including creatine kinase and lactate dehydrogenase, when compared to the control group (**Fig.2J**).

### Deletion of myocardial IGF2R alters transcriptional profiles in response to I/R

RNA sequencing (RNA-seq) was performed to analyze the transcriptome of cKO and Ctrl hearts at baseline and after 12-hour preservation and transplantation (I/R). Principal component analysis of the normalized datasets showed that the transcriptional profiles of the uninjured cKO and Ctrl hearts were similar, while the cKO hearts exhibited distinct expression patterns compared to the other samples following I/R (**Fig.3A**). To gain a deeper understanding of the underlying mechanism of IGF2R, we analyzed the differentially expressed genes in cKO and Ctrl hearts subjected to the I/R condition.

**Fig.3.**
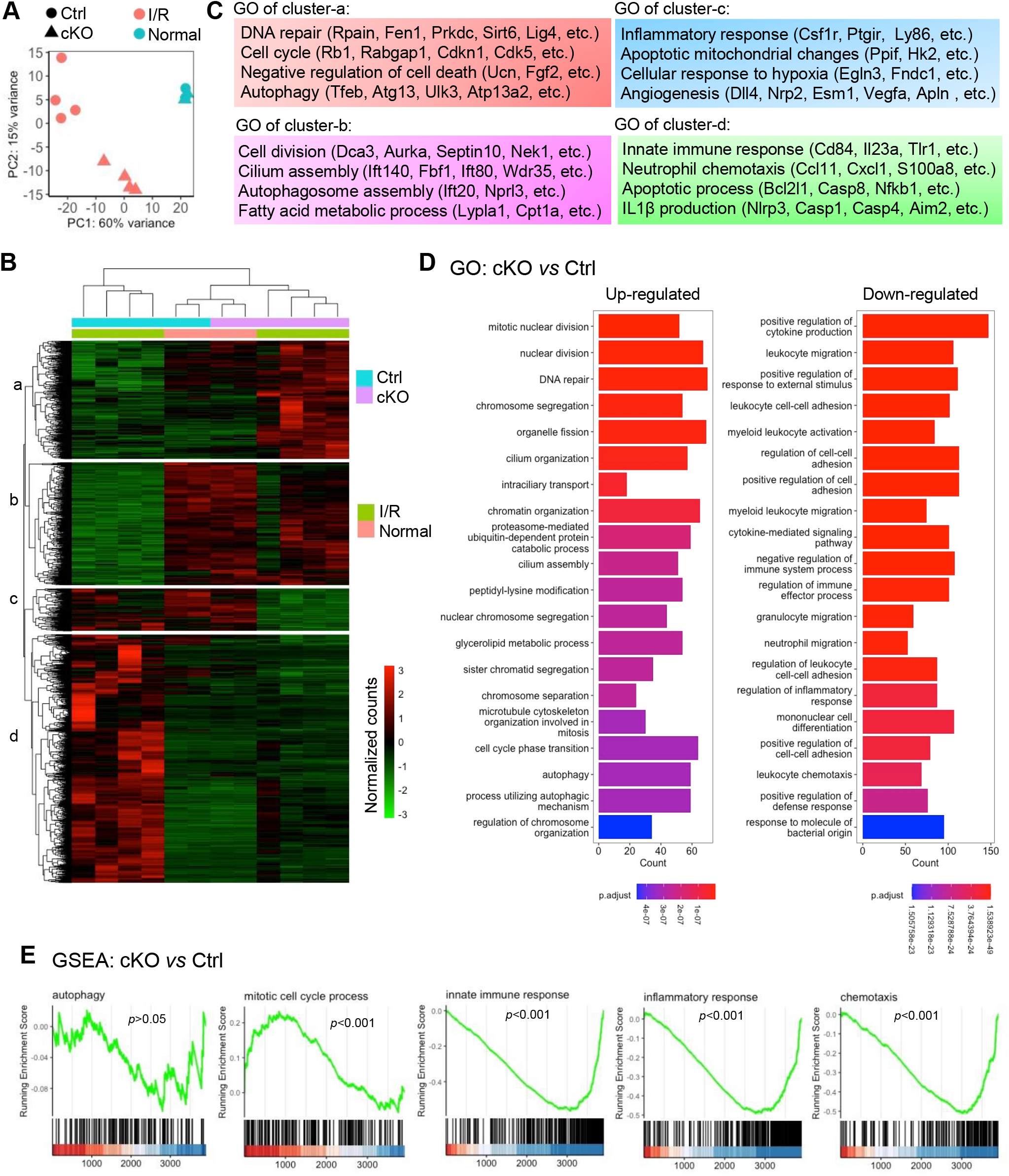
Transcriptional signatures changed by IGF2R-cKO. (**A**) Principal component (PC) analysis of the global gene profiling in the control or cKO heart at normal or I/R (12-hour cold HTK and transplant reperfusion for 12 hours) conditions. (**B-C**) Heatmap and hierarchical clustering analysis of differentially expressed genes in cKO heart as compared to the control. The examples of different gene clusters are shown with their functional annotations (GO). (**D**) Top enriched GO terms of upregulated or downregulated genes in the cKO heart compared with the control. (**E**) GSEA analysis of various GO terms related to the upregulated or downregulated genes in the cKO heart compared with the control.

We identified 3,909 significant genes (*padj*<0.05) differentially expressed in cKO and Ctrl hearts following I/R, and used hierarchical clustering analysis to group these genes into distinct clusters (**Fig.3B**). After I/R, clusters a and b showed up-regulated gene expression in cKO hearts, while clusters c and d exhibited up-regulated gene expression in Ctrl hearts. At baseline, both cKO and Ctrl hearts displayed similar patterns of up-regulated gene expression in clusters b and c, and down-regulated gene expression in clusters a and d. Gene ontology (GO) analysis was used to identify the over-representation of GO terms (biological processes) in the gene lists of different clusters (**Fig.3C**). Genes involved in DNA repair, cell cycle regulation, and autophagy were predominantly represented in clusters a and b, while clusters c and d were enriched for genes related to inflammatory response, innate immune response, and apoptotic process. The top 20 GO terms showed that the cKO of IGF2R significantly attenuated innate immune responses, including cytokine production, leukocyte migration, and cell adhesion, while promoting cell cycle progression and self-healing processes. (**Fig.3D**).

We also performed Gene Set Enrichment Analysis (GSEA) on a ranked list of significant genes to identify enriched gene sets and biological pathways associated with the observed gene expression changes (**Fig.3E**). The results of the gene set distribution and enrichment score using GSEA confirmed that the Ctrl hearts were enriched for GO terms related to inflammatory response, innate immune response, and chemotaxis, whereas the cKO hearts showed enrichment for the mitotic cell cycle process except for autophagy. The cKO of IGF2R resulted in the activation of genes involved in cell growth pathways, which supports the established concept that IGF2R functions as a growth-inhibitory gene.^21, 25^.

### Identification of IGF2R-associated proteins from reperfused myocardium

Although the innate immune response is known to play a critical role in the pathogenesis of myocardial I/R injury,^26, 27^ the precise mechanism by which IGF2R participates in this process is still unknown. Therefore, we attempted to identify the IGF2R targets by proteomic analysis of lysed tissues obtained from heart grafts using pull-down assays on day 1 post-I/R (**Fig.4A**). To this end, we selectively excised protein bands from the control sample, which were missing from the cKO sample at equivalent molecular weights, and performed mass spectrometry analysis. The lists of identified proteins were compared with the RNA-seq results of the down-regulated (log2fc<-1, padj<0.05) or up-regulated (log2fc>1, padj<0.05) genes (**Fig.4B**). Notably, the consistent down-regulation of *Actb, Ngp, Chil3 (Chi3l3), S100a9*, and *Ltf* gene in cKO heart tissue at both transcriptomic and translational levels highlighted their potential role as therapeutic targets of IGF2R. The single-cell RNA-seq dataset (Tabula Muris)^24^ also reveals that these genes are expressed in granulocytes, granulocytopoietic cells, and monocytes in bone marrow, but not in the native heart (**Fig.4C**). At 4 weeks post I/R injury, the Ctrl heart exhibited a substantial increase in cell infiltration with visible nuclei, while the cKO heart demonstrated a significant reduction in cell infiltration (**Fig.4D**). Furthermore, Masson’s trichrome staining showed that the fibrosis area in the cKO heart was smaller than that in the Ctrl heart (**Fig.4E**).

**Fig.4.**
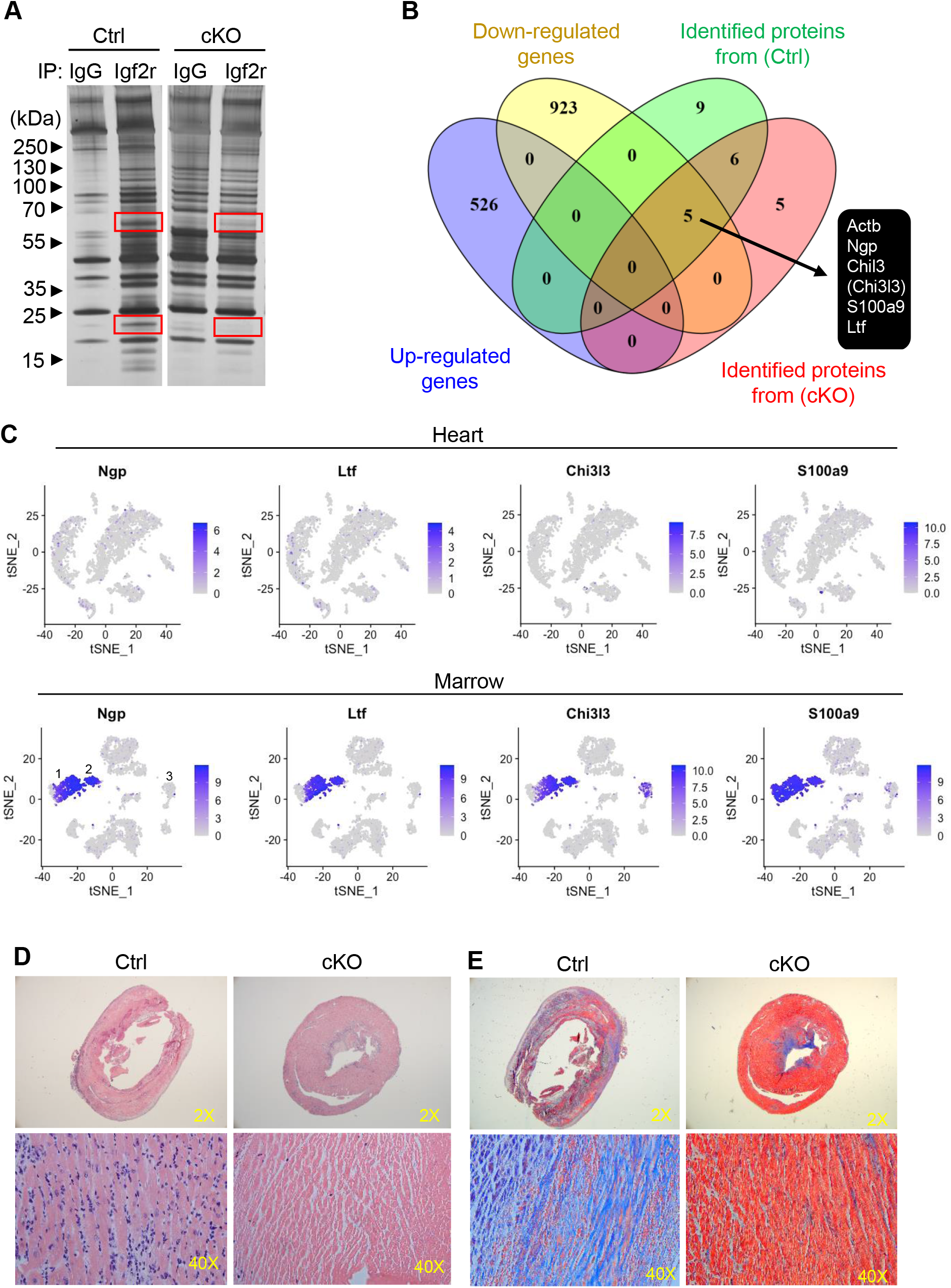
IGF2R binds to granulocyte-derived inflammatory factors. (A) Representative silver staining image of immunoblotting of IgG or anti-IGF2R antibody pulled-down proteins from the lysed control or cKO hearts. Red rectangles labeling the bands for mass spectrometry analysis. (**B**) Venn’s diagram comparing the lists of genes upregulated or downregulated in cKO hearts and identified peptides or proteins from the selected bands from (A). Five factors identified from the integrated analysis are listed. (**C**) Single-cell profiling of Ngp, Chil3 (Chi3l3), S100a9, or Ltf expression in normal mouse hearts (GSM2967050) or bone marrow (GSM2967055). Cell phenotype annotation: 1-granulocytes; 2-granulocytopoietic cell; 3-monocyte. (**D-E**) Representative hematoxylin and eosin staining (D) or Masson’s trichrome staining (E) of the control or cKO heart grafts after I/R (12-hour cold HTK and transplant reperfusion for 4 weeks).

### Hypoxia-induced activation of IGF2R leads to apoptotic cell death

IGF2R was expressed in H9C2 cardiomyoblasts, and *in vitro* model of hypoxia-reoxygenation was used to mimic I/R injury. As compared to the normoxic condition, the IGF2R protein level was increased after hypoxia for 3 hours but slightly decreased after reoxygenation (**Fig.5A-B**). It is known that IGF2R shuttles constantly between intracellular organelles (∼90% of the total IGF2R protein) and the cell surface membrane.^9, 25^ Labeling with wheat germ agglutinin indicated that IGF2R was primarily localized in the intracellular organelles of H9C2 cells (**Fig.5C**). Interestingly, under hypoxia, the intracellular IGF2R appeared to diffuse from the perinuclear region toward the plasma membrane, resulting in a dispersed cytoplasmic distribution. Following hypoxia-reoxygenation, the immunoblot analysis of protein fractions revealed that IGF2R was highly expressed in the intracellular membrane fraction, indicating membrane attachment rather than cytoplasmic solubility (**Fig.5D**).

A doxycycline (Dox)-inducible CRISPR-interference (CRISPRi) system (pHAGE-TRE-dCas9-KRAB)^28^ was established to determine whether the increased IGF2R protein level was due to its promoter transcription activation (**Fig.5E**). The IGF2R protein expression in H9C2 cells was decreased by Dox-induced CRISPRi under normal or hypoxia-reoxygenation conditions (**Fig.5F**). Hypoxia caused a significant increase in the level of cleaved caspase3 (active form), which was attenuated by Dox treatment. Although the cleaved caspase3 level progressively decreased after reoxygenation (to the normoxic level), CRISPRi of IGF2R did not affect cell death during this period.

**Fig.5.**
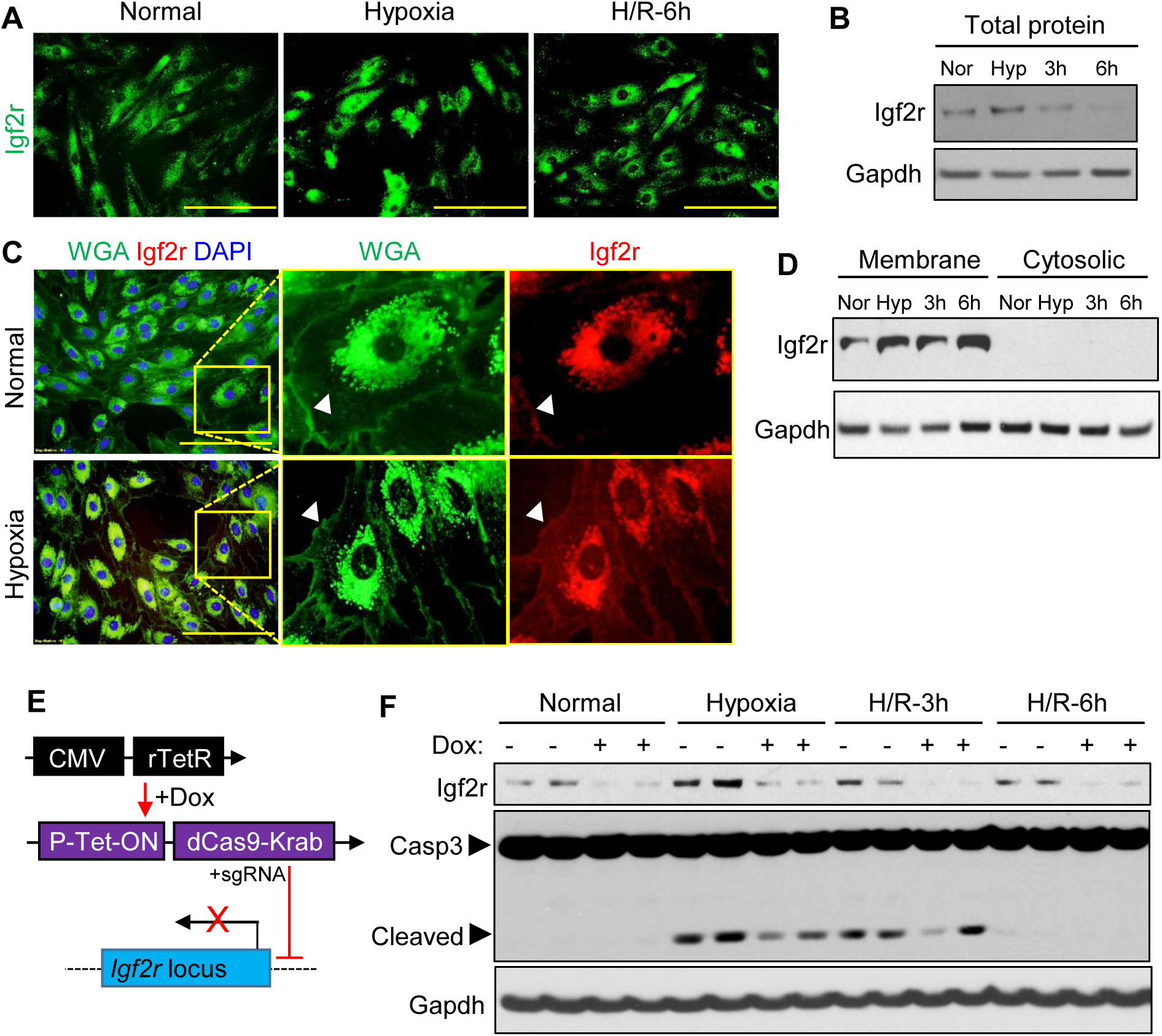
Inhibition of IGF2R expression reduces cell death under hypoxia. (**A**) Immunostaining of IGF2R in H9C2 cardiomyoblasts before or after hypoxia-reoxygenation. Scale bar, 50μm. (**B**) Immunoblotting of IGF2R in total cell proteins from H9C2 myoblasts treated with or without hypoxia-reoxygenation. (**C**) Immunostaining of IGF2R with co-staining of wheat germ agglutinin (WGA) in H9C2 cells before or after hypoxia. Arrowhead indicates the cell plasma membrane. Scale bar, 50μm. (**D**) Immunoblotting of IGF2R after subcellular (membrane or cytosolic) fractions from H9C2 cells before or after hypoxia and reoxygenation. (**E**): H9C2 myoblasts were transduced with the doxycycline (Dox) inducible CRISPRi targeting IGF2R’s promoter region. P-Tet-ON: Pomoter with tetracycline-inducible activation element. (**F**): Immunoblotting of IGF2R and caspase3 in H9C2 cells exposure to hypoxia/reoxygenation.

## DISCUSSION

Although IGF2R is known to be multifaceted, its role in the myocardial I/R response remains poorly characterized. Here, we investigated the role of IGF2R in different models of myocardial I/R injuries. We observed the dynamic expression of IGF2R during myocardial I/R and found that cardiac-specific depletion of IGF2R improved heart dysfunction and reduced cell death. Mechanistic studies revealed that IGF2R plays a role in the inflammatory response and fibrosis by interacting with immune cell-derived proteins. Our findings suggest that IGF2R could be an important therapeutic target for I/R injury and warrant further exploration of its function in cardiac myocytes.

Our results showed that IGF2R expression was upregulated in the ischemic heart but returned to baseline levels following reperfusion. The dynamic expression of IGF2R was also observed in renal macrophages during the I/R processes.^29^ IGF2R is expressed widely in both cardiac and non-cardiac cells, but whether its expression is consistent or variable across different cell types in response to injuries remains to be determined. To this end, single-cell RNA sequencing can be used to characterize the expression of IGF2R across different cell types and heart disease models in future studies. Further research is also needed to fully elucidate the potential transcriptional or epigenetic mechanisms (such as DNA methylation, histone modifications, microRNAs, long non-coding RNAs, and various transcription factors)^30-34^ controlling IGF2R expression in the context of I/R.

Our loss-of-function studies provided novel evidence that myocardial IGF2R played a role in the immune responses following reperfusion or heart transplantation. Depleting cardiac IGF2R expression resulted in a reduction of cell infiltration and fibrosis in the heart tissue after reperfusion injury. The integrated analysis of the transcriptome and proteome highlighted Ngp, Chil3, S100a9, and Ltf as potential targets of IGF2R. Given the selective expression of these genes in bone marrow granulocytes but not in normal cardiac cells, we reasoned that IGF2R could be a key player in regulating the interplay between cardiomyocytes and granulocytes after myocardial damage. Immune cell infiltration is a common response to stress signals in the myocardium and is observed across various types of myocardial injuries.^35^ Neutrophils and other granulocytes can rapidly and massively infiltrate the site of injury and release reactive oxygen species, proteolytic enzymes, and other cytotoxic molecules that can cause further damage to the myocardium and contribute to the development of cardiac fibrosis.^36, 37^ Ngp, Chil3, S100a9, and Ltf are among several proteins secreted by granulocytes that have been implicated in a variety of pathological processes underlying heart disease, including inflammation, immune response, and fibrosis.^38-42^ Therefore, evidence suggests that IGF2R-mediated protein capture may lead to activation of inflammation pathways in cardiomyocytes and subsequent impairment of heart function following I/R injury. However, further investigation is needed to elucidate the precise mechanisms involved.

Furthermore, our findings indicated that myoblasts subjected to hypoxia experienced an increase in IGF2R expression, which was reversed upon reoxygenation. This pattern was consistent with the expression trend of IGF2R observed in heart tissue during the I/R processes. It is possible that the regulation of IGF2R expression during hypoxia and reoxygenation is mediated by different transcription factors or signaling pathways. However, previous studies investigating the impact of hypoxia on the expression of the IGF2R gene have yielded conflicting results. Some studies have shown increased expression of IGF2R during hypoxia in myoblast cells,^17, 18^ while others have found that hypoxia did not affect IGF2R expression in ectoplacental cones.^43^ Therefore, the effects of hypoxia on IGF2R expression appear to be cell type-specific and involve multiple signaling pathways. Aside from its overall gene expression level, the functionality of IGF2R is also dependent on its intracellular localization.^44^ The results of our study demonstrated that hypoxia enhanced the intracellular trafficking of IGF2R from the perinuclear region to the plasma membrane, and this effect was primarily attributed to membrane attachment rather than cytoplasmic solubility.

To assess the therapeutic implications of specifically targeting IGF2R expression, we utilized CRISPRi complex through the Dox-controlled transactivator system to block the IGF2R promoter region in myoblasts. By enabling targeted gene regulation, the CRISPRi technique offers remarkable specificity, versatility, and reversibility, and presents a promising approach for treating a diverse range of diseases.^45, 46^ IGF2R expression was significantly eliminated by CRISPRi which was found to attenuate cell death under hypoxic conditions, yet the molecular pathways involved in this protective effect remain to be determined. Future studies using I/R animal models will be also needed to determine the potential therapeutic effect of this approach.

In summary, this study investigates the expression and function of IGF2R in various models of myocardial I/R injury. We observed dynamic changes in IGF2R expression and intracellular distribution in response to ischemic or hypoxic conditions. Deletion of IGF2R resulted in improved heart function and reduced cell death, indicating its involvement in myocardial I/R injury. RNA sequencing analysis revealed that IGF2R deletion altered transcriptional profiles in response to I/R, attenuating innate immune responses while promoting cell cycle progression and self-healing processes. Proteomic analysis identified granulocyte-specific factors as potential targets of IGF2R binding. Taken together, these findings suggest that myocardial IGF2R plays an essential role in I/R injury and that its deletion may offer a cardioprotective effect against inflammation or fibrosis. This study contributes to our understanding of the pathogenesis of myocardial I/R injury and identifies IGF2R as a potential therapeutic target for ischemic heart disease.

## ACKNOWLEDGMENTS

This work was supported by the American Heart Association award 22TPA968781 (J.L.) and the National Institutes of Health (NIH) grants including R01HL143490 and R01HL157456 (Y.W.). We would like to thank Dr. Randy Jirtle and Dr. David Skaar (North Carolina State University) for providing the *Igf2r*^*fl/fl*^ mice.

## DECLARATION OF INTERESTS

The authors declare no competing interests.

## SUPPLEMENTAL METHODS

All research protocols conformed to the Guidelines for the Care and Use of Laboratory Animals published by the National Institutes of Health (National Academies Press, eighth edition, 2011). All animal use protocols and experiments conducted in this study were approved and oversight by the University of Cincinnati Animal Care and Use Committee. Detailed methods are available in the supplemental information.

## Notes

### Competing Interest Statement

The authors have declared no competing interest.

### Summary of Updates

Figure number revised

